# Novel, highly divergent clones in *Listeria monocytogenes* serotype 4b in North America: Sublineages 782 and 1039, members of the hypervirulent clonal complex 2

**DOI:** 10.64898/2026.08.21.744909

**Authors:** Phillip Brown, Zuzana Kucerova, Philippe Pérot, Asmaa Sadat, James H. Jackson, Driss Elhanafi, Enzo Gadin, Marc Lecuit, Sophia Kathariou

## Abstract

*Listeria monocytogenes* is a Gram-positive bacterial foodborne pathogen responsible for the severe illness listeriosis. Of the 14 *L. monocytogenes* serotypes, serotype 4b is a major contributor to human listeriosis and encompasses all four leading hypervirulent clonal complexes (CCs), including the ancient, ubiquitous CC2. CC2 is globally dominated by sublineage (SL) 2, responsible for most human CC2-associated cases. Here we describe two other CC2 SLs, SLs 782 and 1039. These SLs are newly recognized, having been reported only since 2002, and to date are encountered exclusively in North America. Phylogenetic analysis revealed that they are strikingly divergent from each other as well as from SL2. SL782 and SL1039 have been implicated in human listeriosis and have also been repeatedly isolated from surface water and wildlife in North America, with several of these environmental strains exhibiting high genomic similarity (≤7 core genome allelic mismatches) to strains from human listeriosis. They share an unusual resistance profile towards a panel of *Listeria* wide-host-range-phages and exhibit several distinct lineage-specific traits. Specifically, SL782 universally lacks a gene otherwise unique to and conserved in serotype 4b and harbors the *Listeria* pathogenicity island LIPI-4, while SL1039 harbors LIPI-3 and is almost always resistant to tetracycline, harboring the novel Tn*916*-like transposon Tn*916*.*1039*. These and other traits may have driven clonal emergence of SL782 and SL1039, potentially via adaptations in natural ecosystems.

**IMPORTANCE:** *Listeria monocytogenes* causes the severe foodborne illness listeriosis and the hypervirulent clonal complex (CC) 2 is a major contributor to human disease. Most CC2 strains belong to the long-recognized, ubiquitous sublineage (SL) 2. Surprisingly, the two other leading SLs within CC2, SL782 and 1039, have been isolated exclusively from North America, and only subsequently to 2002. They differ remarkably from each other and other CC2 strains and exhibit unusual genomic and phenotypic traits. They have been repeatedly isolated from watersheds and wildlife, with such environmental isolates exhibiting high genomic similarity to those from human listeriosis. The findings are important in identifying novel, unexpected features in the composition and evolution of a major, hypervirulent *L. monocytogenes* CC. They will also serve as platforms for further studies to further elucidate the reservoirs, emergence, persistence, and dissemination of novel *Listeria* sublineages and their roles in the continuum between natural ecosystems and human disease.

## INTRODUCTION

*Listeria monocytogenes* is a Gram-positive, foodborne, facultative intracellular pathogen and the causative agent of the severe illness listeriosis. Immunosuppression, advanced age, and pregnancy increase the risk for listeriosis. Clinical outcomes of human listeriosis include septicemia, meningitis, encephalitis and stillbirths, with high mortality rates (approx. 20-30%) (1–3). *L. monocytogenes* is ubiquitous in nature and is repeatedly isolated from food and food processing environments (FPE) (1). FPEs are critical in the contamination of listeriosis-implicated food products (1,4). However, surveillance of previously under-explored habitats has identified novel potential reservoirs, e.g., surface waters and wildlife (5–8).

Most cases of human listeriosis belong to two genomic lineages, lineage I (serotypes 1/2b, 4b and 3b) and II (serotypes 1/2a, 1/2c, 3a and 3c) (1). A small proportion of serotype 4b strains, primarily of animal origin, belong to a third lineage, lineage III, uncommonly encountered in human illness (1). Of the 14 recognized serotypes of *L. monocytogenes*, three, i.e., 1/2a, 1/2b and 4b, account for most cases of human listeriosis, with serotype 4b of lineage I making especially significant contributions (9). Multilocus sequence typing (MLST)-based subtyping has revealed major clonal complexes (CCs) that include strains of closely-related sequence types (STs) based on the seven-locus MLST scheme (10). Within a given CC, closely-related strains with <150 allelic mismatches belong to the same sublineage (SL), and sublineages differ by 1,000-1,400 mismatches from each other (11).

Four *L. monocytogenes* CCs (CC1, CC2, CC4 and CC6) are considered hypervirulent, being significantly more frequent in human disease than expected based on incidence in foods, and with relatively high capacity to cause invasive listeriosis via breach of the blood-brain and blood-placenta barriers, thus leading to meningitis or encephalitis and stillbirths, respectively (12). These four hypervirulent CCs are all of serotype 4b (12). CC1, CC2, and CC4 are ubiquitous, ancient clones of *L. monocytogenes* (10), implicated in most early foodborne outbreaks in Europe and North America starting with the late 1970s (10,13). In contrast, CC6 was first recognized with the 1998-1999 hot dog outbreak in the United States (USA) but has since disseminated globally as evidenced by its involvement in 2017 in the largest listeriosis outbreak known to date, in South Africa (10,14,15).

CC1, CC2 and CC6 constitute the largest hypervirulent CCs and, for reasons that remain to be elucidated, they are either dominated or exclusively represented by a single, ubiquitously-encountered sublineage, specifically SL1, SL2, and SL6 in the case of CC1, CC2, and CC6, respectively (11). Analysis of *L. monocytogenes* genomes from diverse sources revealed that all 178 CC1 genomes were SL1, and similarly all 166 CC6 genomes were SL6, while 94 of the 97 analyzed CC2 genomes were SL2 (11). The remaining three CC2 genomes included two of SL781 and one of SL782, with all three strains being derived from the United States (11). The dominance of a single SL in CC1, CC2 and CC6 is rendered even more remarkable by the finding that each of the dominant SLs is itself vastly dominated by a single ST. Thus, ST1 accounted for 95% of the SL1 isolates with available ST designations, while 92 % of the SL2 genomes were ST2, and 99% of the SL6 genomes were ST6 (11).

Considering the pronounced dominance of a single ST in these clones, we became surprised to notice repeated recent isolations in the United States of strains with ST782, belonging to CC2 and previously represented by a single strain, of SL782 (11). ST782 was encountered on several occasions in 2012-2016 during a watershed surveillance in Salinas, California, USA (7), as well as among human clinical isolates in the United States obtained in 2013-2018 (22). Even more surprising was the repeated discovery of ST1039, first identified as a novel ST of CC2 in human clinical isolates in the United States (23). Intriguingly, ST1039 strains were also isolated on multiple occasions in 2014-2016 during surveillance of wildlife (black bears, *Ursus americanus*) in the United States (6).

The unexpected discovery of multiple strains with ST782 and ST1039 within CC2 raises a number of questions. It is currently not clear whether all available ST782 strains belong to the same sublineage (SL782), reported previously with a single strain of this ST (11). It is also unclear whether the different ST1039 strains constitute a separate sublineage, or simply a novel ST in CC2. If they constitute a new sublineage, it would be important to determine its phylogenetic placement in the context of the other known CC2 sublineages. Lastly, it is worthy of note that previously-reported strains with ST782 and ST1039 were all derived from the United States (6,11,23) and had certain unusual traits. For instance, SL782 strains lacked *LMOf2365_1900*, a gene otherwise conserved in serotype 4b, and thus yielded the serotype 1/2b profile via a multiplex PCR assay commonly utilized for molecular assignment of serotype designations (22). On the other hand, the wildlife-derived ST1039 strains harbored a novel Tn*916*-like transposon, Tn*916.1039* (24). The extent to which these unusual traits are conserved within the respective sublineages remains to be determined.

To address these knowledge gaps, in this work we investigated the genome sequences, unique features and origin of all currently-available ST782 and ST1039 strains. This analysis is critically needed in the efforts to elucidate the evolution, global distribution and unique genomic and phenotypic traits of these newly-identified and intriguing STs of the hypervirulent clone CC2.

## MATERIALS AND METHODS

### Bacterial strains, growth conditions, bacteriophage resistance and tetracycline resistance determinations

The strains investigated in this study are listed in Table S1. Additional metadata and specific phenotypic and genomic traits of a subset of these strains (selected SL2 strains from Table S1, and all available strains of the non-SL2 sublineages within CC2, specifically 781, 782, 1039, 1787 and 2377) are listed in Table 1. *L. monocytogenes* was grown in brain heart infusion (BHI) (Becton, Dickinson & Co., Sparks, MD, USA) or BHI with 1.2% agar (BHIA) overnight at 37°C unless otherwise specified. Bacteriophage resistance assays were conducted at 25°C as described (25) using *Listeria* phages A511, P100, 20422-1 and 805405-1 (26–28). Tetracycline minimum inhibitory concentrations (MIC) were determined using MIC strips (Liofilchem, Roseto degli Abruzzi, Teramo, Italy) as per manufacturer instructions and following incubation at 37°C for 24 and 48 hr, in at least two independent trials.

**Table 1.**
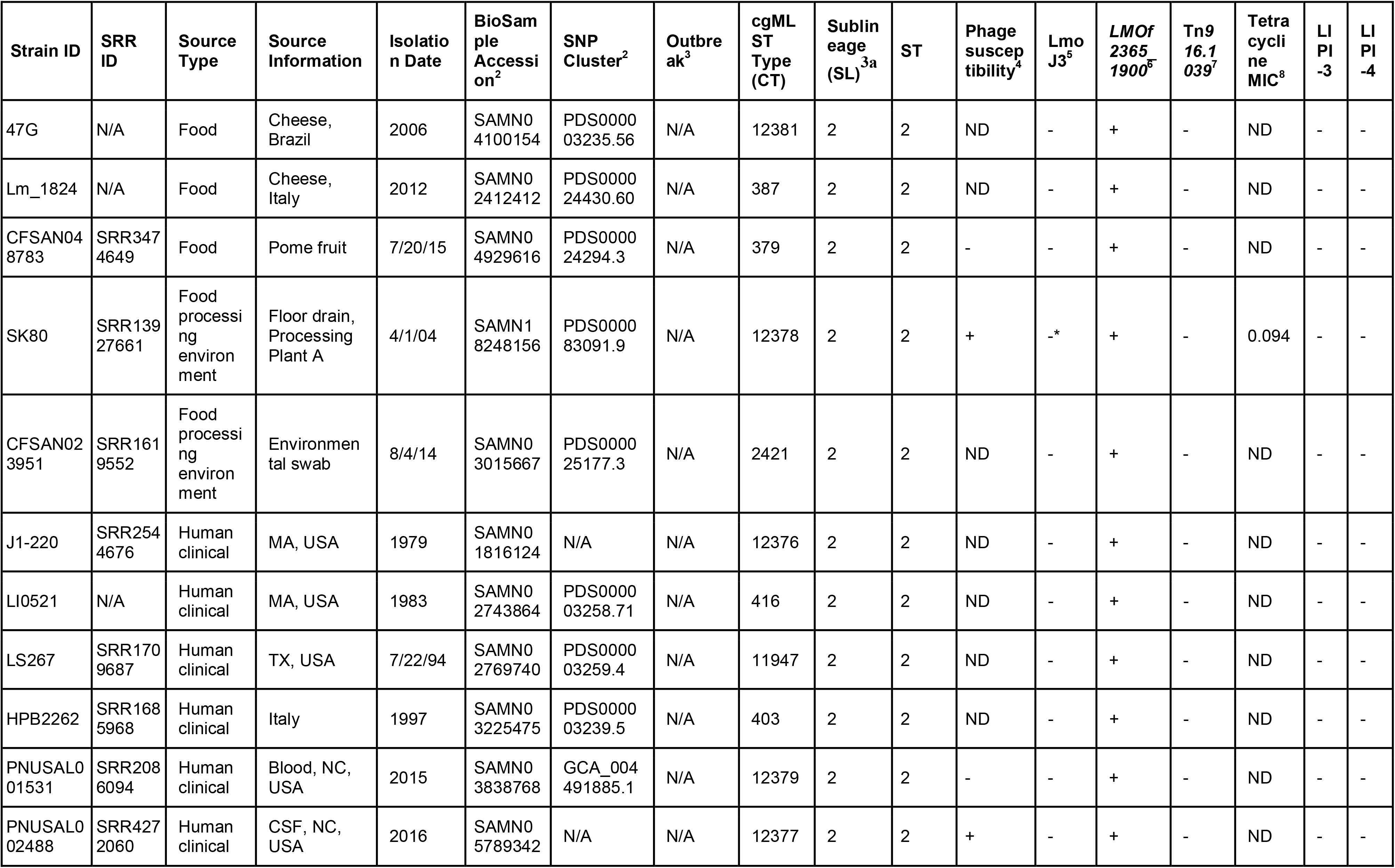
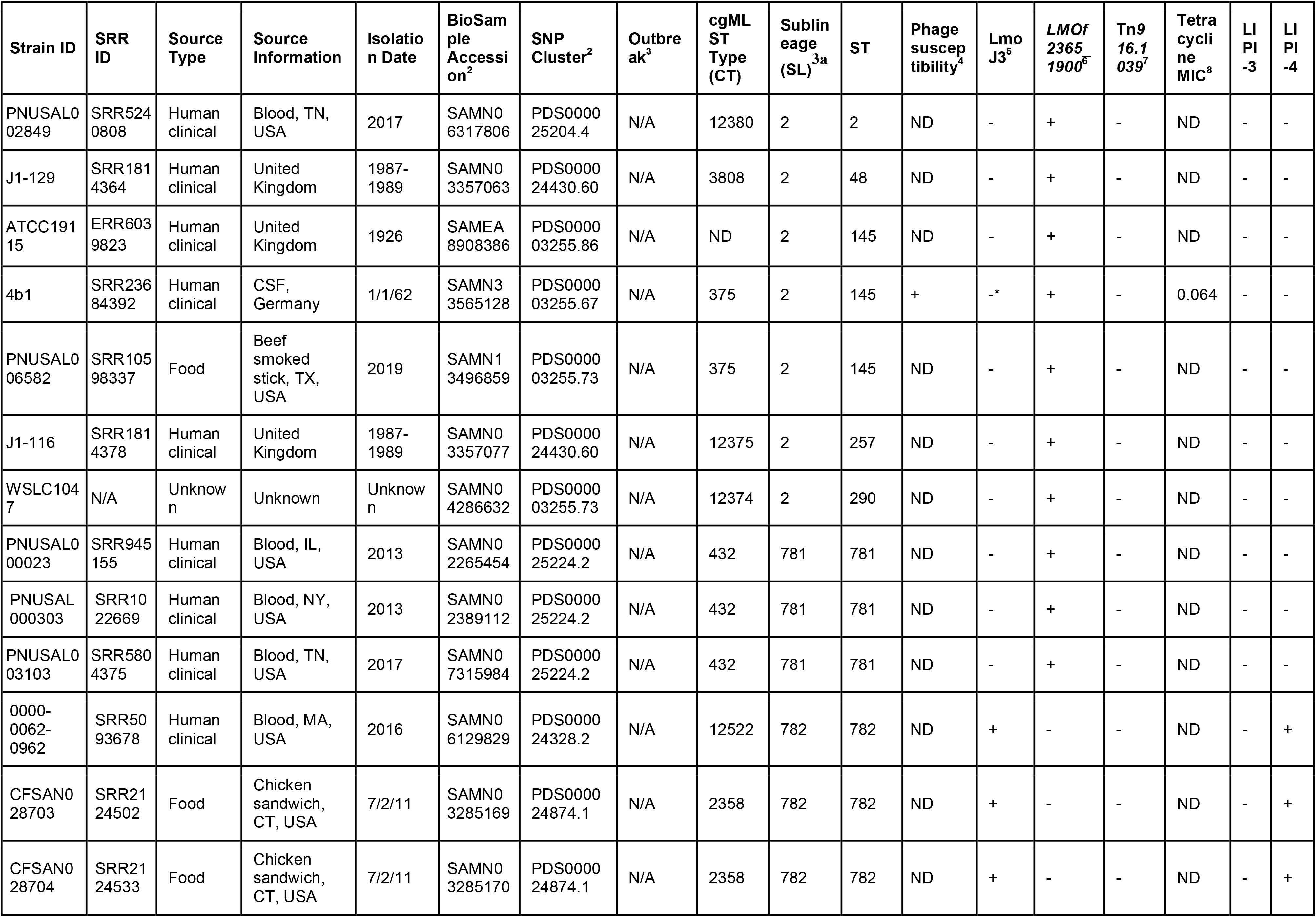
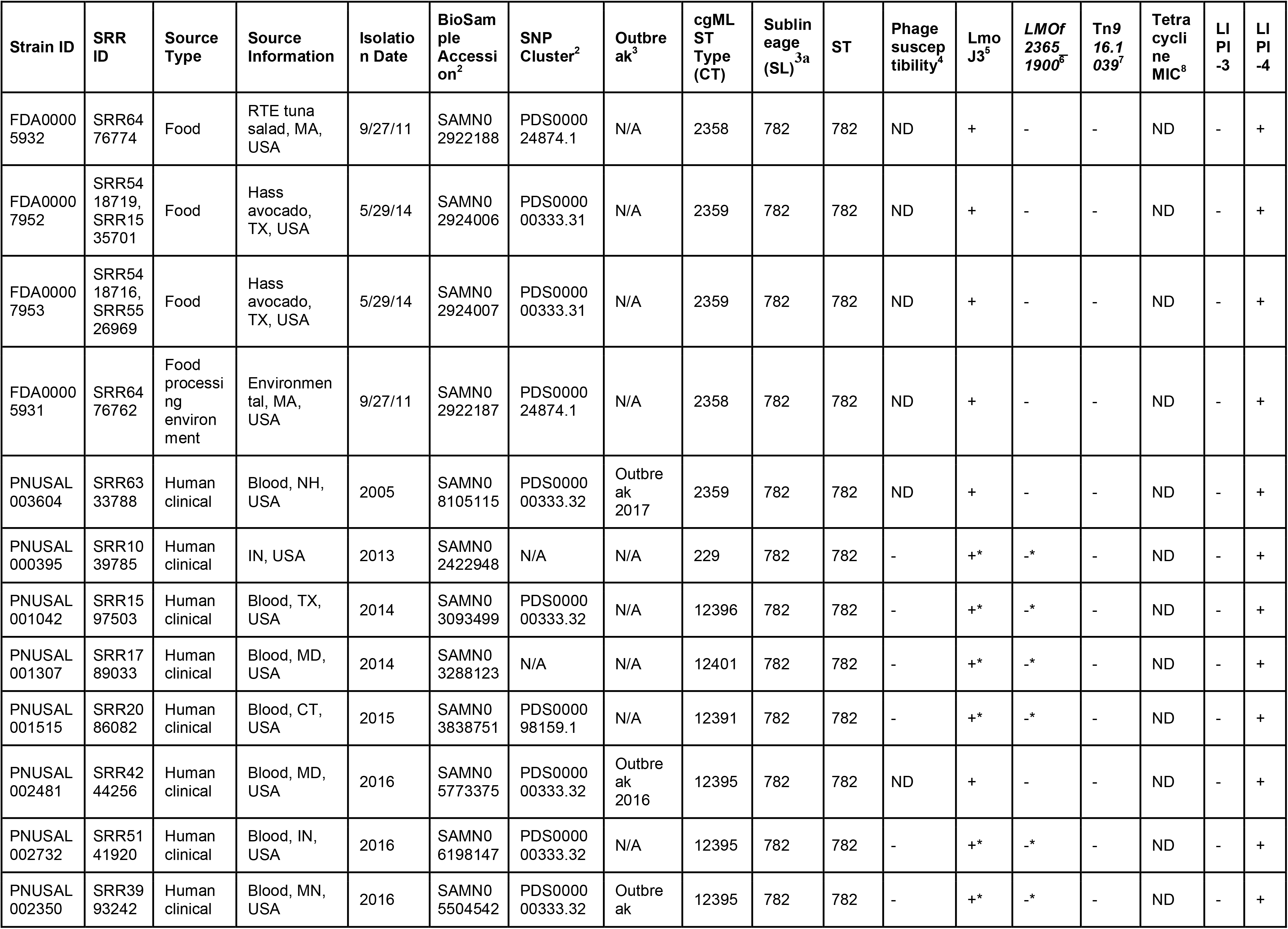
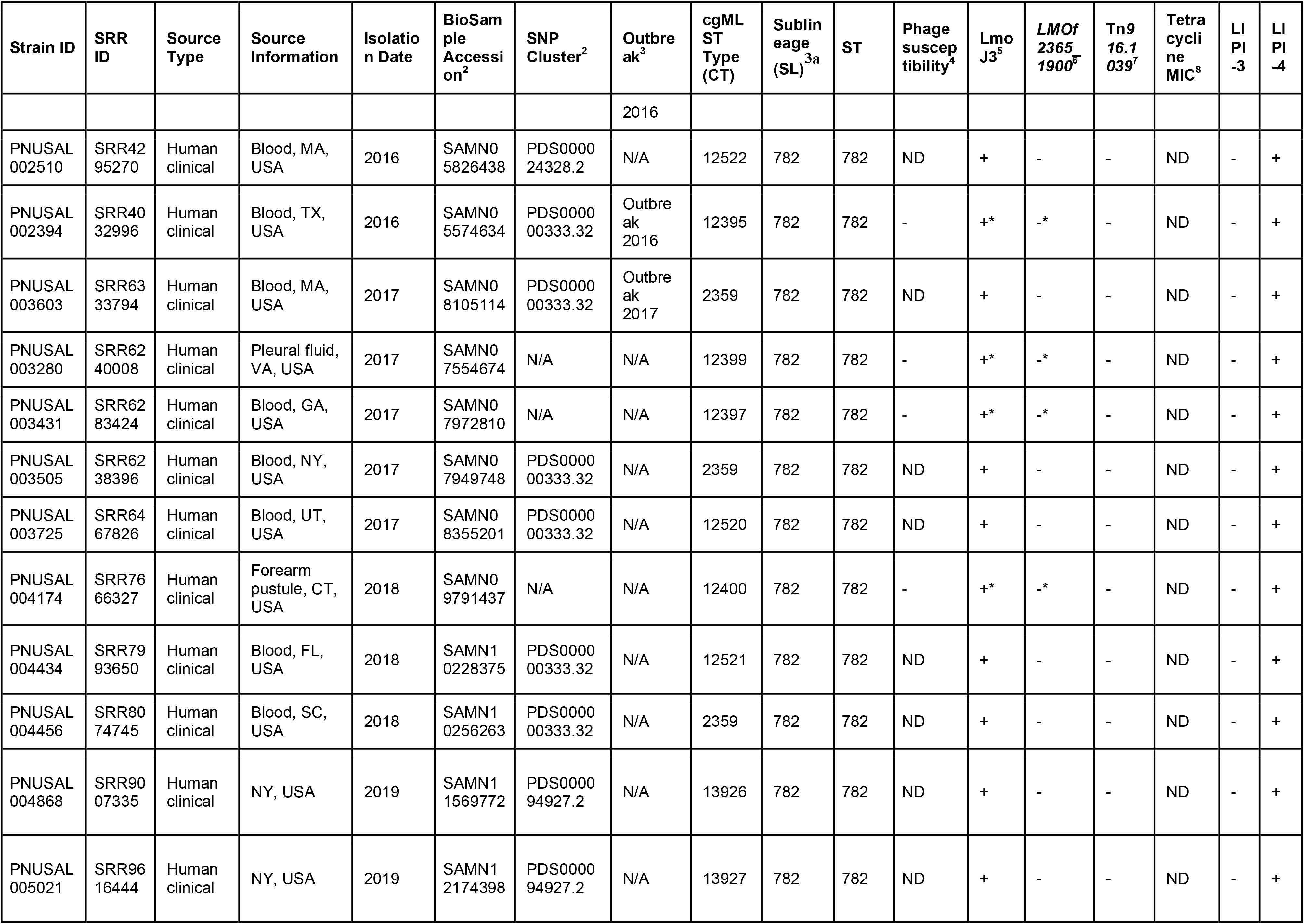
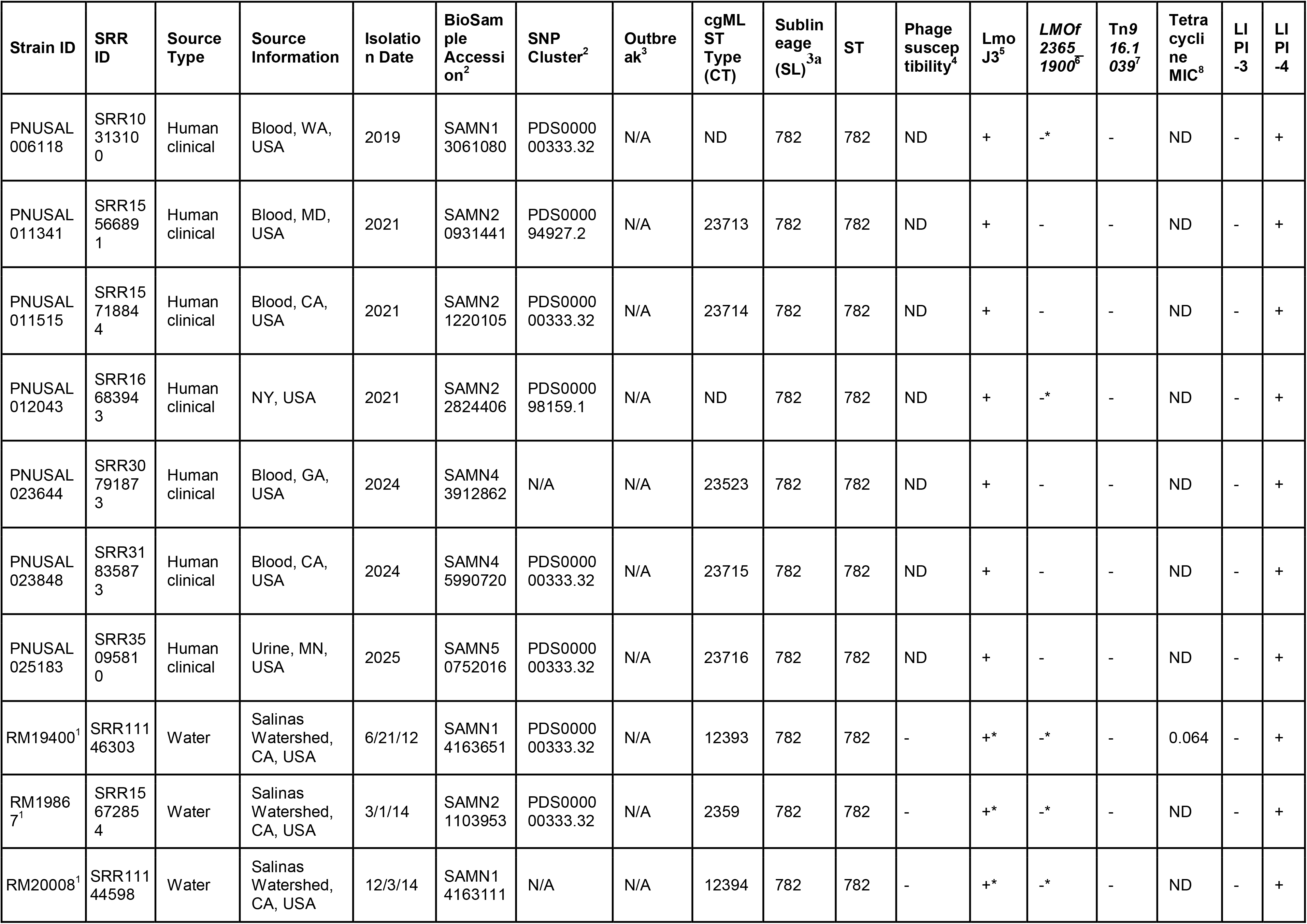
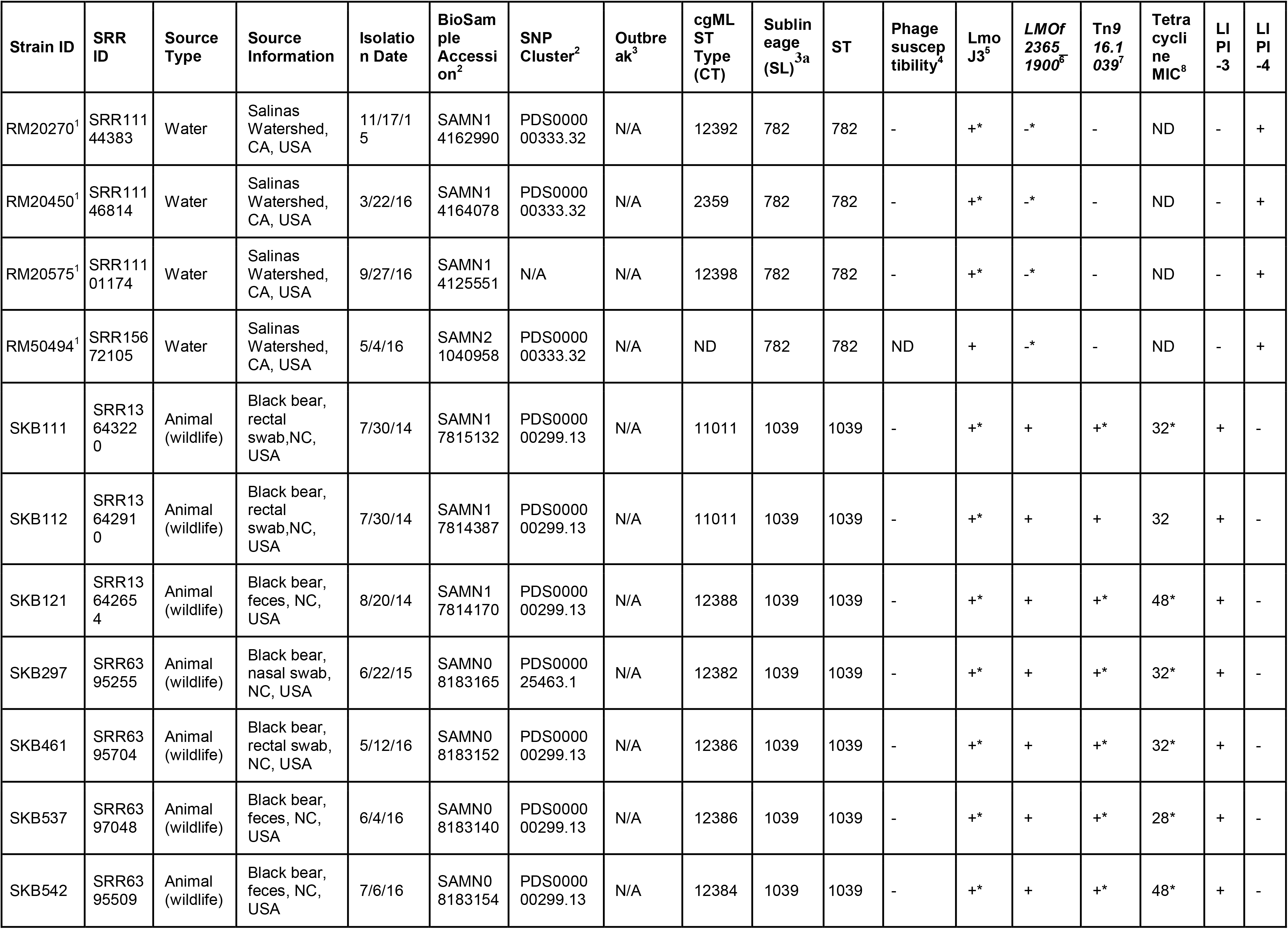
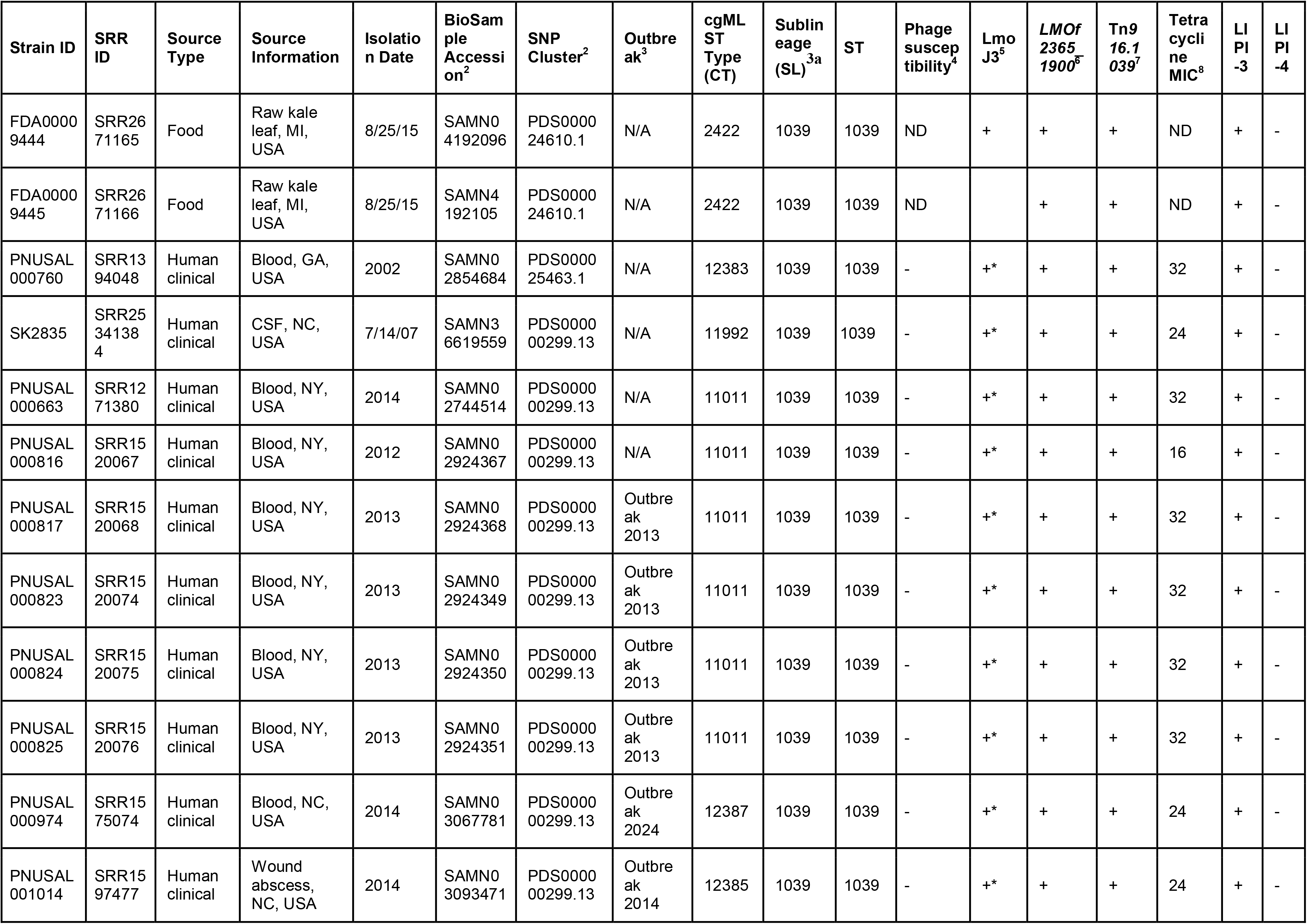
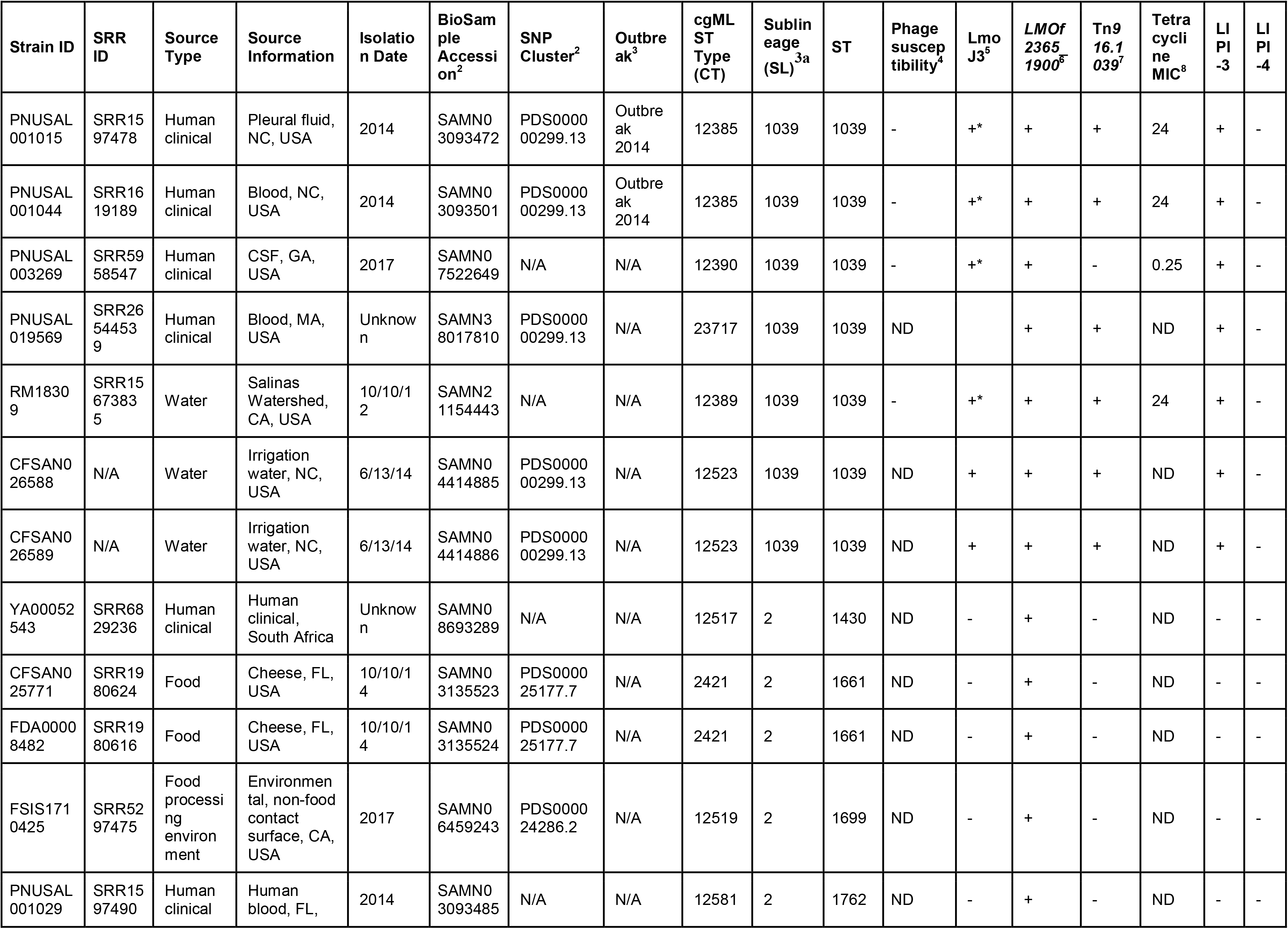
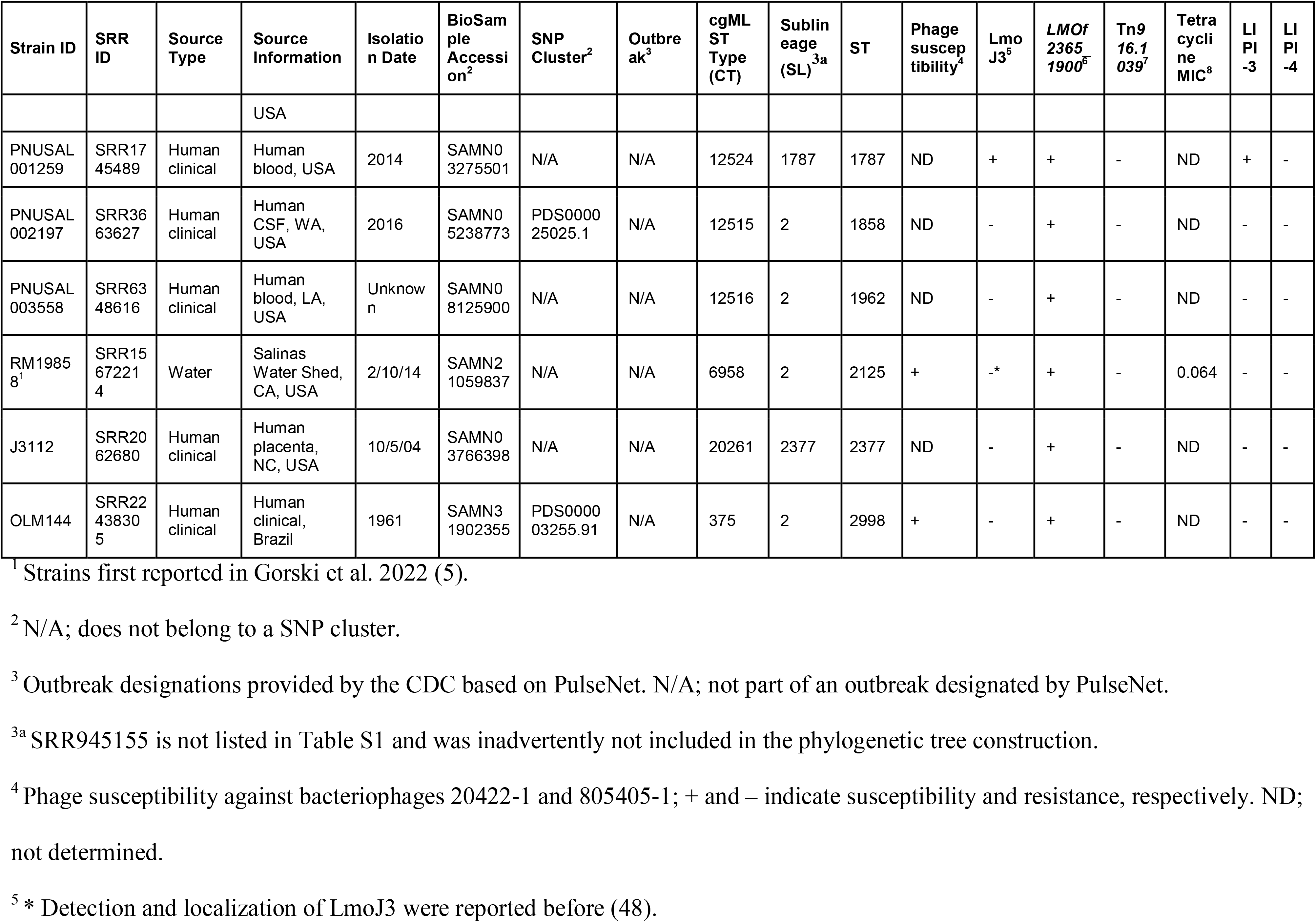
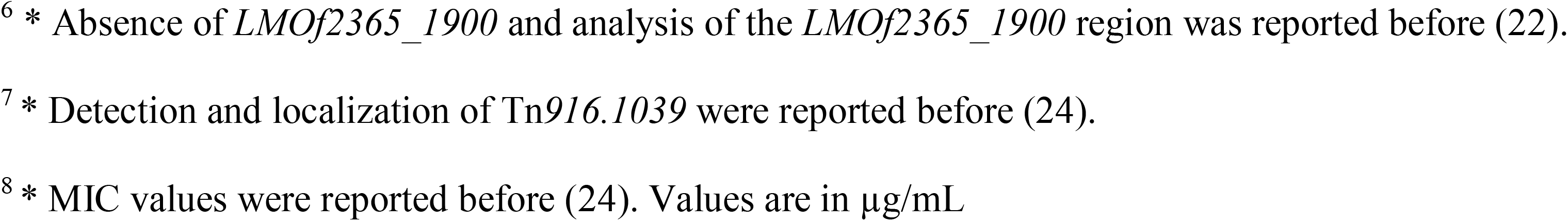
Clonal complex (CC) 2 *Listeria monocytogenes* strains used in this study and their relevant features.

| Strain ID | SRR ID | Source Type | Source Information | Isolation Date | BioSample Accession<sup>2</sup> | SNP Cluster<sup>2</sup> | Outbreak<sup>3</sup> | cgMLST Type (CT) | Sublineage (SL)<sup>3a</sup> | ST | Phage susceptibility<sup>4</sup> | LmoJ3<sup>5</sup> | *LMOf2365_1900*<sup>6</sup> | Tn*916.1039*<sup>7</sup> | Tetracycline MIC<sup>8</sup> | LIPI-3 | LIPI-4 |
|---|---|---|---|---|---|---|---|---|---|---|---|---|---|---|---|---|---|
| 47G | N/A | Food | Cheese, Brazil | 2006 | SAMN04100154 | PDS000003235.56 | N/A | 12381 | 2 | 2 | ND | - | + | - | ND | - | - |
| Lm_1824 | N/A | Food | Cheese, Italy | 2012 | SAMN02412412 | PDS000024430.60 | N/A | 387 | 2 | 2 | ND | - | + | - | ND | - | - |
| CFSAN048783 | SRR3474649 | Food | Pome fruit | 7/20/15 | SAMN04929616 | PDS000024294.3 | N/A | 379 | 2 | 2 | - | - | + | - | ND | - | - |
| SK80 | SRR13927661 | Food processing environment | Floor drain, Processing Plant A | 4/1/04 | SAMN18248156 | PDS000083091.9 | N/A | 12378 | 2 | 2 | + | -* | + | - | 0.094 | - | - |
| CFSAN023951 | SRR1619552 | Food processing environment | Environmental swab | 8/4/14 | SAMN03015667 | PDS000025177.3 | N/A | 2421 | 2 | 2 | ND | - | + | - | ND | - | - |
| J1-220 | SRR2544676 | Human clinical | MA, USA | 1979 | SAMN01816124 | N/A | N/A | 12376 | 2 | 2 | ND | - | + | - | ND | - | - |
| LI0521 | N/A | Human clinical | MA, USA | 1983 | SAMN02743864 | PDS000003258.71 | N/A | 416 | 2 | 2 | ND | - | + | - | ND | - | - |
| LS267 | SRR1709687 | Human clinical | TX, USA | 7/22/94 | SAMN02769740 | PDS000003259.4 | N/A | 11947 | 2 | 2 | ND | - | + | - | ND | - | - |
| HPB2262 | SRR1685968 | Human clinical | Italy | 1997 | SAMN03225475 | PDS000003239.5 | N/A | 403 | 2 | 2 | ND | - | + | - | ND | - | - |
| PNUSAL001531 | SRR2086094 | Human clinical | Blood, NC, USA | 2015 | SAMN03838768 | GCA_004491885.1 | N/A | 12379 | 2 | 2 | - | - | + | - | ND | - | - |
| PNUSAL002488 | SRR4272060 | Human clinical | CSF, NC, USA | 2016 | SAMN05789342 | N/A | N/A | 12377 | 2 | 2 | + | - | + | - | ND | - | - |
| PNUSAL002849 | SRR5240808 | Human clinical | Blood, TN, USA | 2017 | SAMN06317806 | PDS000025204.4 | N/A | 12380 | 2 | 2 | ND | - | + | - | ND | - | - |
| J1-129 | SRR1814364 | Human clinical | United Kingdom | 1987-1989 | SAMN03357063 | PDS000024430.60 | N/A | 3808 | 2 | 48 | ND | - | + | - | ND | - | - |
| ATCC19115 | ERR6039823 | Human clinical | United Kingdom | 1926 | SAMEA8908386 | PDS000003255.86 | N/A | ND | 2 | 145 | ND | - | + | - | ND | - | - |
| 4b1 | SRR23684392 | Human clinical | CSF, Germany | 1/1/62 | SAMN33565128 | PDS000003255.67 | N/A | 375 | 2 | 145 | + | -* | + | - | 0.064 | - | - |
| PNUSAL006582 | SRR10598337 | Food | Beef smoked stick, TX, USA | 2019 | SAMN13496859 | PDS000003255.73 | N/A | 375 | 2 | 145 | ND | - | + | - | ND | - | - |
| J1-116 | SRR1814378 | Human clinical | United Kingdom | 1987-1989 | SAMN03357077 | PDS000024430.60 | N/A | 12375 | 2 | 257 | ND | - | + | - | ND | - | - |
| WSLC1047 | N/A | Unknown | Unknown | Unknown | SAMN04286632 | PDS000003255.73 | N/A | 12374 | 2 | 290 | ND | - | + | - | ND | - | - |
| PNUSAL000023 | SRR945155 | Human clinical | Blood, IL, USA | 2013 | SAMN02265454 | PDS000025224.2 | N/A | 432 | 781 | 781 | ND | - | + | - | ND | - | - |
| PNUSAL000303 | SRR1022669 | Human clinical | Blood, NY, USA | 2013 | SAMN02389112 | PDS000025224.2 | N/A | 432 | 781 | 781 | ND | - | + | - | ND | - | - |
| PNUSAL003103 | SRR5804375 | Human clinical | Blood, TN, USA | 2017 | SAMN07315984 | PDS000025224.2 | N/A | 432 | 781 | 781 | ND | - | + | - | ND | - | - |
| 0000-0062-0962 | SRR5093678 | Human clinical | Blood, MA, USA | 2016 | SAMN06129829 | PDS000024328.2 | N/A | 12522 | 782 | 782 | ND | + | - | - | ND | - | + |
| CFSAN028703 | SRR2124502 | Food | Chicken sandwich, CT, USA | 7/2/11 | SAMN03285169 | PDS000024874.1 | N/A | 2358 | 782 | 782 | ND | + | - | - | ND | - | + |
| CFSAN028704 | SRR2124533 | Food | Chicken sandwich, CT, USA | 7/2/11 | SAMN03285170 | PDS000024874.1 | N/A | 2358 | 782 | 782 | ND | + | - | - | ND | - | + |
| FDA00005932 | SRR6476774 | Food | RTE tuna salad, MA, USA | 9/27/11 | SAMN02922188 | PDS000024874.1 | N/A | 2358 | 782 | 782 | ND | + | - | - | ND | - | + |
| FDA00007952 | SRR5418719, SRR1535701 | Food | Hass avocado, TX, USA | 5/29/14 | SAMN02924006 | PDS000000333.31 | N/A | 2359 | 782 | 782 | ND | + | - | - | ND | - | + |
| FDA00007953 | SRR5418716, SRR5526969 | Food | Hass avocado, TX, USA | 5/29/14 | SAMN02924007 | PDS000000333.31 | N/A | 2359 | 782 | 782 | ND | + | - | - | ND | - | + |
| FDA00005931 | SRR6476762 | Food processing environment | Environmental, MA, USA | 9/27/11 | SAMN02922187 | PDS000024874.1 | N/A | 2358 | 782 | 782 | ND | + | - | - | ND | - | + |
| PNUSAL003604 | SRR6333788 | Human clinical | Blood, NH, USA | 2005 | SAMN08105115 | PDS000000333.32 | Outbreak 2017 | 2359 | 782 | 782 | ND | + | - | - | ND | - | + |
| PNUSAL000395 | SRR1039785 | Human clinical | IN, USA | 2013 | SAMN02422948 | N/A | N/A | 229 | 782 | 782 | - | +* | -* | - | ND | - | + |
| PNUSAL001042 | SRR1597503 | Human clinical | Blood, TX, USA | 2014 | SAMN03093499 | PDS000000333.32 | N/A | 12396 | 782 | 782 | - | +* | -* | - | ND | - | + |
| PNUSAL001307 | SRR1789033 | Human clinical | Blood, MD, USA | 2014 | SAMN03288123 | N/A | N/A | 12401 | 782 | 782 | - | +* | -* | - | ND | - | + |
| PNUSAL001515 | SRR2086082 | Human clinical | Blood, CT, USA | 2015 | SAMN03838751 | PDS000098159.1 | N/A | 12391 | 782 | 782 | - | +* | -* | - | ND | - | + |
| PNUSAL002481 | SRR4244256 | Human clinical | Blood, MD, USA | 2016 | SAMN05773375 | PDS000000333.32 | Outbreak 2016 | 12395 | 782 | 782 | ND | + | - | - | ND | - | + |
| PNUSAL002732 | SRR5141920 | Human clinical | Blood, IN, USA | 2016 | SAMN06198147 | PDS000000333.32 | N/A | 12395 | 782 | 782 | - | +* | -* | - | ND | - | + |
| PNUSAL002350 | SRR3993242 | Human clinical | Blood, MN, USA | 2016 | SAMN05504542 | PDS000000333.32 | Outbreak 2016 | 12395 | 782 | 782 | - | +* | -* | - | ND | - | + |
| PNUSAL002510 | SRR4295270 | Human clinical | Blood, MA, USA | 2016 | SAMN05826438 | PDS000024328.2 | N/A | 12522 | 782 | 782 | ND | + | - | - | ND | - | + |
| PNUSAL002394 | SRR4032996 | Human clinical | Blood, TX, USA | 2016 | SAMN05574634 | PDS000000333.32 | Outbreak 2016 | 12395 | 782 | 782 | - | +* | -* | - | ND | - | + |
| PNUSAL003603 | SRR6333794 | Human clinical | Blood, MA, USA | 2017 | SAMN08105114 | PDS000000333.32 | Outbreak 2017 | 2359 | 782 | 782 | ND | + | - | - | ND | - | + |
| PNUSAL003280 | SRR6240008 | Human clinical | Pleural fluid, VA, USA | 2017 | SAMN07554674 | N/A | N/A | 12399 | 782 | 782 | - | +* | -* | - | ND | - | + |
| PNUSAL003431 | SRR6283424 | Human clinical | Blood, GA, USA | 2017 | SAMN07972810 | N/A | N/A | 12397 | 782 | 782 | - | +* | -* | - | ND | - | + |
| PNUSAL003505 | SRR6238396 | Human clinical | Blood, NY, USA | 2017 | SAMN07949748 | PDS000000333.32 | N/A | 2359 | 782 | 782 | ND | + | - | - | ND | - | + |
| PNUSAL003725 | SRR6467826 | Human clinical | Blood, UT, USA | 2017 | SAMN08355201 | PDS000000333.32 | N/A | 12520 | 782 | 782 | ND | + | - | - | ND | - | + |
| PNUSAL004174 | SRR7666327 | Human clinical | Forearm pustule, CT, USA | 2018 | SAMN09791437 | N/A | N/A | 12400 | 782 | 782 | - | +* | -* | - | ND | - | + |
| PNUSAL004434 | SRR7993650 | Human clinical | Blood, FL, USA | 2018 | SAMN10228375 | PDS000000333.32 | N/A | 12521 | 782 | 782 | ND | + | - | - | ND | - | + |
| PNUSAL004456 | SRR8074745 | Human clinical | Blood, SC, USA | 2018 | SAMN10256263 | PDS000000333.32 | N/A | 2359 | 782 | 782 | ND | + | - | - | ND | - | + |
| PNUSAL004868 | SRR9007335 | Human clinical | NY, USA | 2019 | SAMN11569772 | PDS000094927.2 | N/A | 13926 | 782 | 782 | ND | + | - | - | ND | - | + |
| PNUSAL005021 | SRR9616444 | Human clinical | NY, USA | 2019 | SAMN12174398 | PDS000094927.2 | N/A | 13927 | 782 | 782 | ND | + | - | - | ND | - | + |
| PNUSAL006118 | SRR10313100 | Human clinical | Blood, WA, USA | 2019 | SAMN13061080 | PDS000000333.32 | N/A | ND | 782 | 782 | ND | + | -* | - | ND | - | + |
| PNUSAL011341 | SRR15566891 | Human clinical | Blood, MD, USA | 2021 | SAMN20931441 | PDS000094927.2 | N/A | 23713 | 782 | 782 | ND | + | - | - | ND | - | + |
| PNUSAL011515 | SRR15718844 | Human clinical | Blood, CA, USA | 2021 | SAMN21220105 | PDS000000333.32 | N/A | 23714 | 782 | 782 | ND | + | - | - | ND | - | + |
| PNUSAL012043 | SRR16683943 | Human clinical | NY, USA | 2021 | SAMN22824406 | PDS000098159.1 | N/A | ND | 782 | 782 | ND | + | -* | - | ND | - | + |
| PNUSAL023644 | SRR30791873 | Human clinical | Blood, GA, USA | 2024 | SAMN43912862 | N/A | N/A | 23523 | 782 | 782 | ND | + | - | - | ND | - | + |
| PNUSAL023848 | SRR31835873 | Human clinical | Blood, CA, USA | 2024 | SAMN45990720 | PDS000000333.32 | N/A | 23715 | 782 | 782 | ND | + | - | - | ND | - | + |
| PNUSAL025183 | SRR35095810 | Human clinical | Urine, MN, USA | 2025 | SAMN50752016 | PDS000000333.32 | N/A | 23716 | 782 | 782 | ND | + | - | - | ND | - | + |
| RM19400<sup>1</sup> | SRR11146303 | Water | Salinas Watershed, CA, USA | 6/21/12 | SAMN14163651 | PDS000000333.32 | N/A | 12393 | 782 | 782 | - | +* | -* | - | 0.064 | - | + |
| RM19867<sup>1</sup> | SRR15672854 | Water | Salinas Watershed, CA, USA | 3/1/14 | SAMN21103953 | PDS000000333.32 | N/A | 2359 | 782 | 782 | - | +* | -* | - | ND | - | + |
| RM20008<sup>1</sup> | SRR11144598 | Water | Salinas Watershed, CA, USA | 12/3/14 | SAMN14163111 | N/A | N/A | 12394 | 782 | 782 | - | +* | -* | - | ND | - | + |
| RM20270<sup>1</sup> | SRR11144383 | Water | Salinas Watershed, CA, USA | 11/17/15 | SAMN14162990 | PDS000000333.32 | N/A | 12392 | 782 | 782 | - | +* | -* | - | ND | - | + |
| RM20450<sup>1</sup> | SRR11146814 | Water | Salinas Watershed, CA, USA | 3/22/16 | SAMN14164078 | PDS000000333.32 | N/A | 2359 | 782 | 782 | - | +* | -* | - | ND | - | + |
| RM20575<sup>1</sup> | SRR11101174 | Water | Salinas Watershed, CA, USA | 9/27/16 | SAMN14125551 | N/A | N/A | 12398 | 782 | 782 | - | +* | -* | - | ND | - | + |
| RM50494<sup>1</sup> | SRR15672105 | Water | Salinas Watershed, CA, USA | 5/4/16 | SAMN21040958 | PDS000000333.32 | N/A | ND | 782 | 782 | ND | + | -* | - | ND | - | + |
| SKB111 | SRR13643220 | Animal (wildlife) | Black bear, rectal swab,NC, USA | 7/30/14 | SAMN17815132 | PDS000000299.13 | N/A | 11011 | 1039 | 1039 | - | +* | + | +* | 32* | + | - |
| SKB112 | SRR13642910 | Animal (wildlife) | Black bear, rectal swab,NC, USA | 7/30/14 | SAMN17814387 | PDS000000299.13 | N/A | 11011 | 1039 | 1039 | - | +* | + | + | 32 | + | - |
| SKB121 | SRR13642654 | Animal (wildlife) | Black bear, feces, NC, USA | 8/20/14 | SAMN17814170 | PDS000000299.13 | N/A | 12388 | 1039 | 1039 | - | +* | + | +* | 48* | + | - |
| SKB297 | SRR6395255 | Animal (wildlife) | Black bear, nasal swab, NC, USA | 6/22/15 | SAMN08183165 | PDS000025463.1 | N/A | 12382 | 1039 | 1039 | - | +* | + | +* | 32* | + | - |
| SKB461 | SRR6395704 | Animal (wildlife) | Black bear, rectal swab, NC, USA | 5/12/16 | SAMN08183152 | PDS000000299.13 | N/A | 12386 | 1039 | 1039 | - | +* | + | +* | 32* | + | - |
| SKB537 | SRR6397048 | Animal (wildlife) | Black bear, feces, NC, USA | 6/4/16 | SAMN08183140 | PDS000000299.13 | N/A | 12386 | 1039 | 1039 | - | +* | + | +* | 28* | + | - |
| SKB542 | SRR6395509 | Animal (wildlife) | Black bear, feces, NC, USA | 7/6/16 | SAMN08183154 | PDS000000299.13 | N/A | 12384 | 1039 | 1039 | - | +* | + | +* | 48* | + | - |
| FDA00009444 | SRR2671165 | Food | Raw kale leaf, MI, USA | 8/25/15 | SAMN04192096 | PDS000024610.1 | N/A | 2422 | 1039 | 1039 | ND | + | + | + | ND | + | - |
| FDA00009445 | SRR2671166 | Food | Raw kale leaf, MI, USA | 8/25/15 | SAMN4192105 | PDS000024610.1 | N/A | 2422 | 1039 | 1039 | ND |  | + | + | ND | + | - |
| PNUSAL000760 | SRR1394048 | Human clinical | Blood, GA, USA | 2002 | SAMN02854684 | PDS000025463.1 | N/A | 12383 | 1039 | 1039 | - | +* | + | + | 32 | + | - |
| SK2835 | SRR25341384 | Human clinical | CSF, NC, USA | 7/14/07 | SAMN36619559 | PDS000000299.13 | N/A | 11992 | 1039 | 1039 | - | +* | + | + | 24 | + | - |
| PNUSAL000663 | SRR1271380 | Human clinical | Blood, NY, USA | 2014 | SAMN02744514 | PDS000000299.13 | N/A | 11011 | 1039 | 1039 | - | +* | + | + | 32 | + | - |
| PNUSAL000816 | SRR1520067 | Human clinical | Blood, NY, USA | 2012 | SAMN02924367 | PDS000000299.13 | N/A | 11011 | 1039 | 1039 | - | +* | + | + | 16 | + | - |
| PNUSAL000817 | SRR1520068 | Human clinical | Blood, NY, USA | 2013 | SAMN02924368 | PDS000000299.13 | Outbreak 2013 | 11011 | 1039 | 1039 | - | +* | + | + | 32 | + | - |
| PNUSAL000823 | SRR1520074 | Human clinical | Blood, NY, USA | 2013 | SAMN02924349 | PDS000000299.13 | Outbreak 2013 | 11011 | 1039 | 1039 | - | +* | + | + | 32 | + | - |
| PNUSAL000824 | SRR1520075 | Human clinical | Blood, NY, USA | 2013 | SAMN02924350 | PDS000000299.13 | Outbreak 2013 | 11011 | 1039 | 1039 | - | +* | + | + | 32 | + | - |
| PNUSAL000825 | SRR1520076 | Human clinical | Blood, NY, USA | 2013 | SAMN02924351 | PDS000000299.13 | Outbreak 2013 | 11011 | 1039 | 1039 | - | +* | + | + | 32 | + | - |
| PNUSAL000974 | SRR1575074 | Human clinical | Blood, NC, USA | 2014 | SAMN03067781 | PDS000000299.13 | Outbreak 2024 | 12387 | 1039 | 1039 | - | +* | + | + | 24 | + | - |
| PNUSAL001014 | SRR1597477 | Human clinical | Wound abscess, NC, USA | 2014 | SAMN03093471 | PDS000000299.13 | Outbreak 2014 | 12385 | 1039 | 1039 | - | +* | + | + | 24 | + | - |
| PNUSAL001015 | SRR1597478 | Human clinical | Pleural fluid, NC, USA | 2014 | SAMN03093472 | PDS000000299.13 | Outbreak 2014 | 12385 | 1039 | 1039 | - | +* | + | + | 24 | + | - |
| PNUSAL001044 | SRR1619189 | Human clinical | Blood, NC, USA | 2014 | SAMN03093501 | PDS000000299.13 | Outbreak 2014 | 12385 | 1039 | 1039 | - | +* | + | + | 24 | + | - |
| PNUSAL003269 | SRR5958547 | Human clinical | CSF, GA, USA | 2017 | SAMN07522649 | N/A | N/A | 12390 | 1039 | 1039 | - | +* | + | - | 0.25 | + | - |
| PNUSAL019569 | SRR26544539 | Human clinical | Blood, MA, USA | Unknown | SAMN38017810 | PDS000000299.13 | N/A | 23717 | 1039 | 1039 | ND |  | + | + | ND | + | - |
| RM18309 | SRR15673835 | Water | Salinas Watershed, CA, USA | 10/10/12 | SAMN21154443 | N/A | N/A | 12389 | 1039 | 1039 | - | +* | + | + | 24 | + | - |
| CFSAN026588 | N/A | Water | Irrigation water, NC, USA | 6/13/14 | SAMN04414885 | PDS000000299.13 | N/A | 12523 | 1039 | 1039 | ND | + | + | + | ND | + | - |
| CFSAN026589 | N/A | Water | Irrigation water, NC, USA | 6/13/14 | SAMN04414886 | PDS000000299.13 | N/A | 12523 | 1039 | 1039 | ND | + | + | + | ND | + | - |
| YA00052543 | SRR6829236 | Human clinical | Human clinical, South Africa | Unknown | SAMN08693289 | N/A | N/A | 12517 | 2 | 1430 | ND | - | + | - | ND | - | - |
| CFSAN025771 | SRR1980624 | Food | Cheese, FL, USA | 10/10/14 | SAMN03135523 | PDS000025177.7 | N/A | 2421 | 2 | 1661 | ND | - | + | - | ND | - | - |
| FDA00008482 | SRR1980616 | Food | Cheese, FL, USA | 10/10/14 | SAMN03135524 | PDS000025177.7 | N/A | 2421 | 2 | 1661 | ND | - | + | - | ND | - | - |
| FSIS1710425 | SRR5297475 | Food processing environment | Environmental, non-food contact surface, CA, USA | 2017 | SAMN06459243 | PDS000024286.2 | N/A | 12519 | 2 | 1699 | ND | - | + | - | ND | - | - |
| PNUSAL001029 | SRR1597490 | Human clinical | Human blood, FL, USA | 2014 | SAMN03093485 | N/A | N/A | 12581 | 2 | 1762 | ND | - | + | - | ND | - | - |
| PNUSAL001259 | SRR1745489 | Human clinical | Human blood, USA | 2014 | SAMN03275501 | N/A | N/A | 12524 | 1787 | 1787 | ND | + | + | - | ND | + | - |
| PNUSAL002197 | SRR3663627 | Human clinical | Human CSF, WA, USA | 2016 | SAMN05238773 | PDS000025025.1 | N/A | 12515 | 2 | 1858 | ND | - | + | - | ND | - | - |
| PNUSAL003558 | SRR6348616 | Human clinical | Human blood, LA, USA | Unknown | SAMN08125900 | N/A | N/A | 12516 | 2 | 1962 | ND | - | + | - | ND | - | - |
| RM19858<sup>1</sup> | SRR15672214 | Water | Salinas Water Shed, CA, USA | 2/10/14 | SAMN21059837 | N/A | N/A | 6958 | 2 | 2125 | + | -* | + | - | 0.064 | - | - |
| J3112 | SRR2062680 | Human clinical | Human placenta, NC, USA | 10/5/04 | SAMN03766398 | N/A | N/A | 20261 | 2377 | 2377 | ND | - | + | - | ND | - | - |
| OLM144 | SRR22438305 | Human clinical | Human clinical, Brazil | 1961 | SAMN31902355 | PDS000003255.91 | N/A | 375 | 2 | 2998 | + | - | + | - | ND | - | - |

<sup>1</sup> Strains first reported in Gorski et al. 2022 (5).
<sup>2</sup> N/A; does not belong to a SNP cluster.
<sup>3</sup> Outbreak designations provided by the CDC based on PulseNet. N/A; not part of an outbreak designated by PulseNet.
<sup>3a</sup> SRR945155 is not listed in Table S1 and was inadvertently not included in the phylogenetic tree construction.
<sup>4</sup> Phage susceptibility against bacteriophages 20422-1 and 805405-1; + and – indicate susceptibility and resistance, respectively. ND; not determined.
<sup>5</sup> \* Detection and localization of LmoJ3 were reported before (48).
<sup>6</sup> \* Absence of *LMOf2365\_1900* and analysis of the *LMOf2365\_1900* region was reported before (22).
<sup>7</sup> \* Detection and localization of Tn*916.1039* were reported before (24).
<sup>8</sup> \* MIC values were reported before (24). Values are in µg/mL

### Core genome sequence and other bioinformatic analysis

Nucleotide BLAST analyses used the National Center for Biotechnology Information (https://blast.ncbi.nlm.nih.gov/Blast.cgi) (29) and the Bacterial and Viral Bioinformatics Resource Center (BV-BRC; https://www.bv-brc.org) (30). Whole genome sequence data for CC2 strains in Fig. S1 were uploaded to the BIGSdb hosted by Institut Pasteur (https://bigsdb.pasteur.fr/listeria/) (31) to determine core genome MLST (cgMLST) profiles using default parameters and excluding incomplete alleles. These cgMLST profiles were used to construct cgMLST-based phylogenetic trees with BioNumerics version 8.1 (http://www.biomerieux.com/). A subset of CC2 strains were screened via BV-BRC and BIGSdb for *L. monocytogenes* pathogenicity islands, the type II restriction modification (RM) system LmoJ3 and other genomic features described previously (12,32).

### Whole genome phylogenetic analyses

High-quality genomes of *L. monocytogenes* CC2 (n=794) were selected from BIGSdb-Lm (https://bigsdb.pasteur.fr/listeria/). These isolates originated from humans (n=427; 53.8%), other animals (n=12; 1.5%), food (n=221; 27.8%), FPEs (n=77; 9.7%), the natural environment (n=43; 5.4%) or unknown sources (n=12; 1.5%). Isolates were from 41 countries and six continents between 1975 and 2024, with one strain, ATCC19115 (Li 2), isolated in 1926 (Table S1). Raw sequencing reads were downloaded from the European Nucleotide Archive (https://www.ebi.ac.uk/ena/browser/home). Raw reads were trimmed to remove adapter sequences and non-confident bases (minimum read length of 30 bases, minimum quality Phred score of 20) using fqCleanER v23.12 (https://gitlab.pasteur.fr/GIPhy/fqCleanER). Alignments based on wgSNP were built from trimmed reads using the Snippy v.4.6.0 pipeline with default parameters (https://github.com/tseemann/snippy). The CC2 genome FSL J1-225 (ScottA; accession no. GCA_000212455.1) was used as reference in read mapping, resulting in an alignment of 3.02 Mb. Gubbins v.3.2.0 (33) was used to detect recombination regions in the whole-genome alignment using default parameters, and alignment regions positive for recombination were removed from the original alignments, resulting in a whole-genome alignment of 2.67 Mb. *SNP-sites* v2.5.1 (34) was used to keep columns containing exclusively ACGT in the alignment, resulting in an alignment of 1.48 Mb. Maximum likelihood phylogeny was obtained from the recombination-purged alignment using IQ-TREE v.2.4.0 (35) under the determined best-fit nucleotide substitution model (K3Pu+F+I), as determined by ModelFinder (36), and ultrafast bootstrapping of 1000 replicates (37). Trees were visualized and annotated with iTol v.7.0 (38).

## RESULTS

### L. monocytogenes strains with ST782 and ST1039 constitute two highly divergent sublineages (SL782 and SL1039) of the hypervirulent CC2

Analysis of the cgMLST data of the ST782 strains previously reported from watershed samples and human clinical cases in the United States (8, 23) revealed that they all were members of SL782 (https://bigsdb.pasteur.fr/listeria/) along with the previously reported human isolate, also from the United States (11). We were able to identify several additional ST782 human isolates via analysis of the CDC *Listeria* database and the NCBI Pathogen Detection Pipeline. These were also determined to be members of SL782. SL782 consists exclusively of strains with ST782 (Table 1).

Analysis of the cgMLST profiles of all available strains of ST1039, including those from wildlife and from human listeriosis (Table 1) revealed that they were strikingly different (>1,000 cgMLST allele differences) not only from SL782 but also from SL2, the dominant sublineage within CC2. They exhibited similarly pronounced divergence from the other CC2 sublineages listed in pubMLST: SL781, previously identified in two strains (11), as well as SL1787 and 2377, encountered once each (https://bigsdb.pasteur.fr/listeria/). These findings support the assignment of ST1039 strains to a novel sublineage, SL1039, which consists exclusively of ST1039 strains (Table 1, Table S2). Thus, CC2 currently includes six sublineages: SLs 2, 781, 782, 1039, 1787, and 2377.

As discussed earlier, SL2 is by far the predominant sublineage in CC2 (11) accounting for 97% of the approx. 4,100 CC2 isolates in PubMLST BIGSdb. Of the five other CC2 sublineages, three, i.e., SLs 781 (n=3), 1787 (n=1) and 2377 (n=1) are extremely uncommon (Table S1). Thus, SL782 (n=30) and SL1039 (n=24) are the only CC2 sublineages other than SL2 encountered among multiple isolates and isolated repeatedly from both clinical and environmental sources (Table S1). All CC2 sublineages other than SL2 have been isolated relatively recently. SL781, SL 1787 and SL2377 were isolated in 2004-2017 while the first known isolations of SL782 and SL1039 were in 2005 and 2002, respectively (Table 1).

Phylogenetic analysis suggested two major clades within CC2: One with just SL2 and the other with all remaining SLs. In the latter, SL782 partitioned in one cluster together with SL1787, while SL1039 partitioned in a separate cluster, together with SL781 and SL2377 (Fig. 1). Little intraclonal diversity was noted within SL782 or SL1039 (Fig. 1), with <100 intra-clone cgMLST allele differences (Table S1). In each sublineage only one strain slightly diverged from the others: PNUSAL000395 (SL782) and PNUSAL003269 (SL1039) (Fig. 1; Table S1).

**Figure 1.**
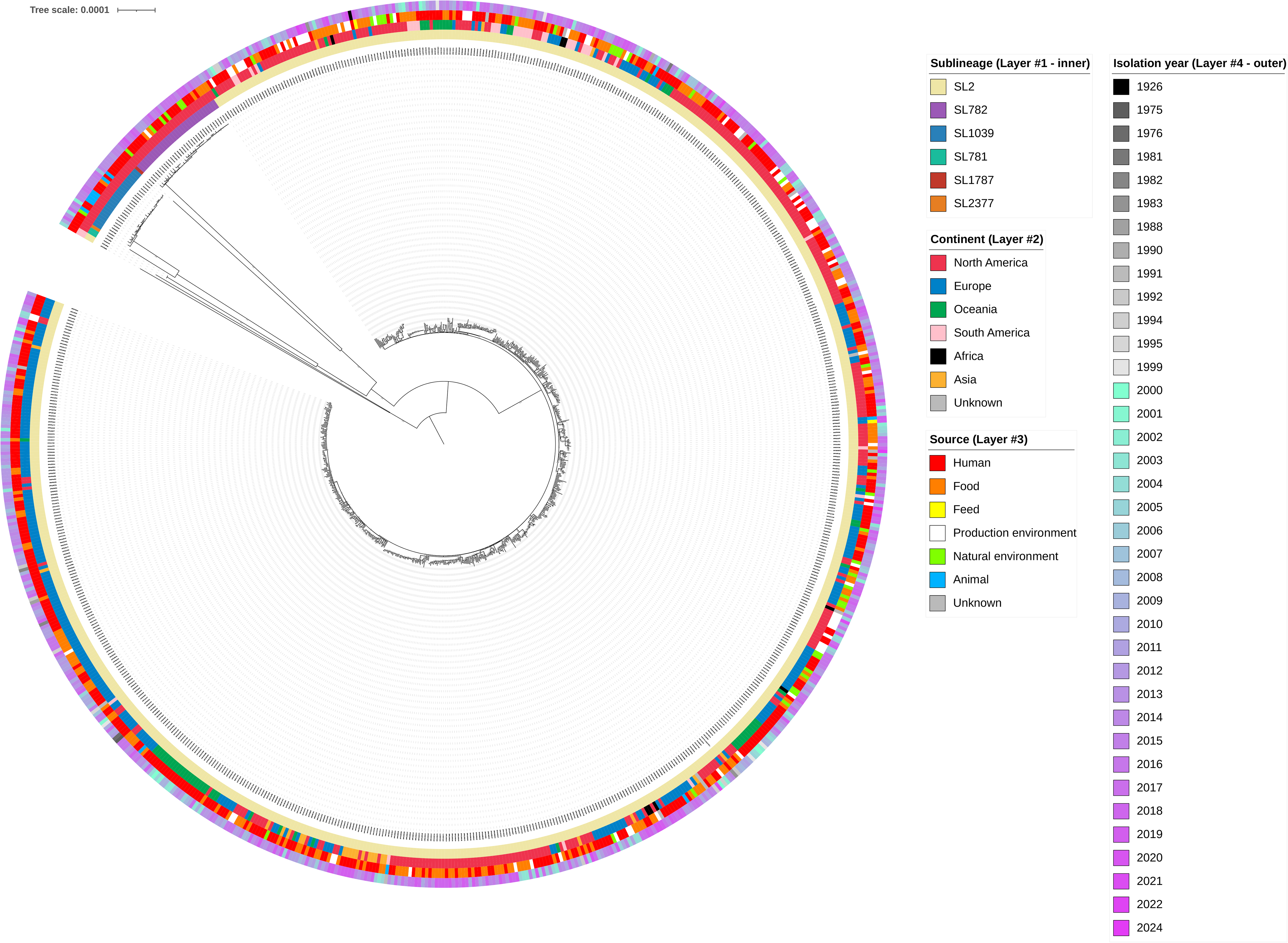
Midpoint-rooted maximum likelihood phylogenetic tree of 794 *Lm*-CC2 genomes based on a recombination-purged whole genome alignment of 1.48 Mb. The four rings, from inner to outer, indicate sublineage, world region, source type and isolation year, respectively. Bootstrap values (1000 replicates, ultrafast bootstrapping) are indicated on the branches in light grey. Tree was visualized in iTOL v.7.0.

Interestingly, SL2 was also remarkably homogeneous, despite its global abundance and highly diverse representation among different regions and time periods; most strains clustered together, with the notable exception of two relatively recent (2004 and 2010) human clinical strains from South America (Fig. 1).

We analyzed the cgMLST profiles of SL782 and SL1039 genomes for cgMLST types (CTs), i.e., clusters of cgMLST profiles with no more than seven allelic mismatches (11). This analysis revealed clusters with shared CT designations that included strains from human listeriosis as well as environmental isolates. Notably, CT2359 included SL782 strains from human cases (n=4), food (n=2, avocado) and water (n=2) while CT11011 consisted of SL1039 isolates derived from human clinical cases (n=6) and wildlife (n=2) (Fig. S1; Table S2). CTs consisting only of human isolates were also identified within each SL, such as CT12395 (n=4) and CT12385 (n=3) in SL782 and SL1039, respectively (Table 1; Fig. S1; Table S3).

### SL782 and SL1039 have been recovered exclusively from North America

The relevant metadata highlight an unusual regional bias. To date, all strains of SL782 and ST1039 are from North America, specifically the United States (Table 1, Table S3). An unusual geographic distribution was furthermore noticed for human clinical strains of SL1039 within the United States. These strains derived only from North Carolina, Georgia, Massachusetts, and New York, all in the eastern USA. In contrast, clinical SL782 strains were from various states representing diverse regions, i.e., midwestern, southeastern, southwestern, western, and northeastern USA (Fig. 2; Table 1).

**Figure 2.**
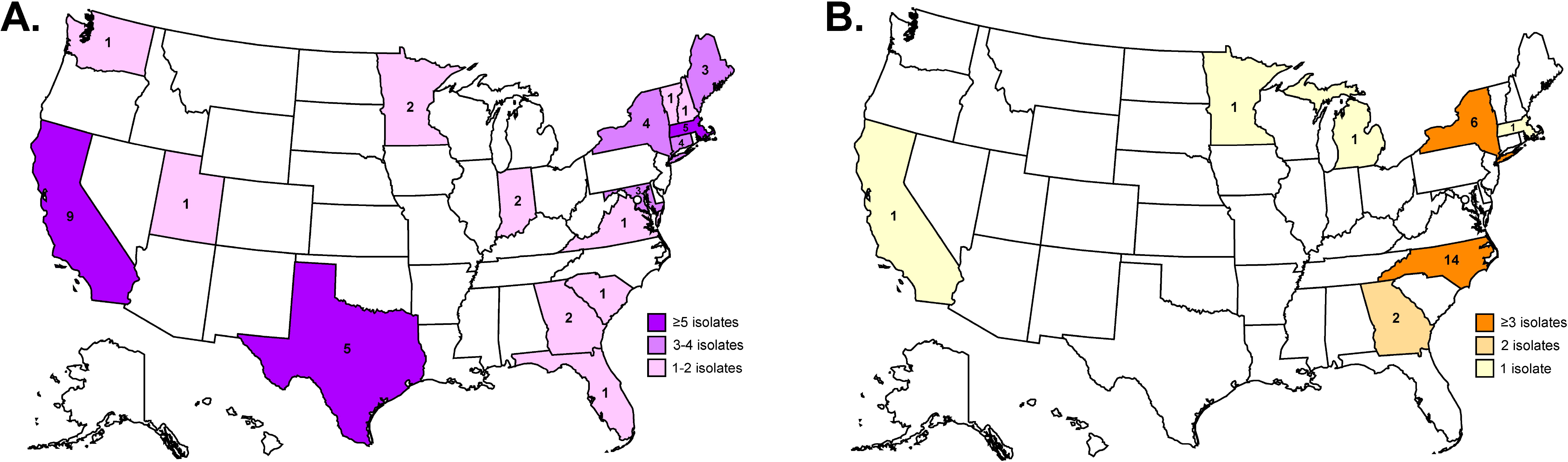
Distribution of *Listeria monocytogenes* (**A**) SL782 and (**B**) SL1039 isolates reported in this study. The number of isolates for each state is indicated with the respective color keys (SL782 - purple, SL1039 - orange) s. **A**. Number of SL782 isolates per state: California (9), Texas (5), Massachusetts (5), Connecticut (4), New York (4), Maine (3), Maryland (3), Georgia (2), Indiana (2), Minnesota (2), Florida (1), New Hampshire (1), South Carolina (1), Utah (1), Vermont (1), Virginia (1), and Washington (1). **B.** Number of SL1039 isolates per state: North Carolina (14), New York (6), California (1), Georgia (1), Massachusetts (1), Michigan (1), and Minnesota (1).

As indicated above, CC2 includes six sublineages. It is of interest that SL781, SL 1787 and SL2377 were also from the United States, similarly to SL782 and SL1039 (Table 1).

### SL1039 and SL782 harbor Listeria Pathogenicity Island (LIPI)-3 and LIPI-4, respectively

SL782 and SL1039 were positive for LIPI-1, as is typical for *L. monocytogenes*, and were expected to have full-length InlA. Interestingly, LIPI-3 was found in all SL1039 strains while LIPI-4 (previously considered unique to SL4 and related clones) was harbored by all SL782 strains and the one available strain of SL1787 (n=1) (Table 1). None of the tested strains harbored LIPI-2 (the LIPI-1 counterpart of *L. ivanovii*), GI-7 (virulence-associated in serotype 4h), stress response-related islets SSI-1 and SSI-2, sanitizer resistance markers LGI-1 (*ermE*) and *qacA*, or the cadmium resistance genes *cadA3* and *cadC3* (Table 1; Table S3). SL782 and SL1039 lacked the quaternary ammonium disinfectant resistance genes *bcrABC* and Tn*6188*-associated *ermC*; these were only detected in SL2, once each. LGI-2 (cadmium and arsenic resistance) and the cadmium resistance cassettes *cadA1C1* and *cadA2C2* were also encountered only in SL2, and were absent from SL782 or SL1039 (Table 1; Table S3).

### Absence of the serotype 4b-specific gene LMOf2365_1900, and high prevalence of the tetracycline resistance transposon Tn916.1039

All known SL782 strains, regardless of origin, lacked *LMOf2365_1900*, a gene otherwise specific for, and unique to, lineage I serotype 4b strains (Table 1). *LMOf2365_1900* was detected in all other CC2 sublineages and all other tested lineage I serotype 4b strains (Table 1).

All but one of the clinical SL1039 strains were resistant to tetracycline and harbored Tn*916.1039* with the tetracycline resistance determinant *tet*(M) previously identified in the wildlife-derived SL1039 strains (24). The transposon was in a conserved chromosomal location, between *coaBC* (*lmo1825*, *LMOf2365_1853*) and *rpoZ* (*lmo1826*, *LMOf2365_1854*) (Table 1).

Only one SL1039 strain, PNUSAL003269 from human cerebrospinal fluid, lacked Tn*916.1039* and was also tetracycline susceptible (Table 1). Interestingly, this strain diverged slightly from other SL1039 strains in the cgMLST-based phylogenetic tree (Fig. 1; Fig. S1; Table 1).

Tn*916.1039* was not detected in SL781 or SL2377, which clustered with SL1039 in the phylogenetic tree (Fig. 1). It was also absent from other CC2 strains or other *L. monocytogenes* clonal complexes, appearing to be unique to SL1039.

### SLs 782 and 1039 share unusual bacteriophage susceptibility profiles and harbor the type II restriction modification system LmoJ3

All tested SL782 and 1039 strains exhibited unusual, clonally-associated bacteriophage susceptibility profiles with four wide-host-range *Listeria* phages: they were susceptible to phages A511 and P100, but resistant to phages 20422-1 and 805405-1 (Table 1). Furthermore, all SL782 and SL1039 genomes harbored a highly-conserved (>99.9% identity, 100% coverage) type II RM system, LmoJ3, in a chromosomal hotspot between *LMOf2365_322* (*lmo0301*) and *LMOf2365_330* (*lmo0305*) (Table 1). In CC2, LmoJ3 was only encountered in SL782, SL1039 and SL1787 which, as indicated earlier, clustered with SL782 (Table 1; Fig. 1).

## DISCUSSION

SL782 and SL1039 constitute two of the six CC2 sublineages and, besides SL2, are the only sublineages of CC2 from multiple strains and diverse sources. They are noteworthy in being isolated recently, i.e., since 2002, and only from North America, specifically the United States.

These sublineages were therefore largely absent from previous studies of *L. monocytogenes* population diversity (12,21,39,40). Even though SL2 remains by far the dominant sublineage in CC2 for human listeriosis, SL782 and SL1039 do have consistent representation in human illness, including their apparent involvement in outbreaks of listeriosis (Table 1).

Interestingly, both SL782 and SL1039 have been repeatedly isolated from natural ecosystems. SL782 was encountered several times during a watershed surveillance in California, USA (8); SL1039 was also detected in that surveillance (5) as well as in agricultural waters in North Carolina, USA (Table 1). Intriguingly, SL1039 was also repeatedly isolated from feces, nares and rectal swabs of wild black bears (*Ursus americanus*) in North Carolina, USA (6). Such findings raise the possibility of SL782 and SL1039 reservoirs in natural, previously under-surveyed ecosystems. The striking phylogenetic divergence of SL782 and 1039 from SL2 the other CC2 sublineages (Fig. 1; Fig. S1; Table S1), further supports evolution in specialized reservoirs with relatively limited access to humans and the human food supply.

SL782 and SL1039 exhibit unusual clonal features possibly accompanying their emergence and readily differentiating them from other CC2 strains. Specifically, our data support and expand the previous finding that ST782 (which we have shown constitutes SL782) lacks the otherwise serotype 4b-specific *LMOf2365_1900* (22), raising the possibility that SL782 emergence was accompanied by loss of this gene. To our knowledge, SL782 is the only serotype 4b sublineage lacking *LMOf2365_1900.* Our data also support and expand the previous finding from analysis of wildlife-derived ST1039 strains, which indicated that these strains harbored the novel Tn*916*-like transposon Tn*916.1039* (24). Intriguingly, Tn*916.1039* was not encountered among tetracycline-resistant *L. innocua* strains from the same wildlife (black bear, *Ursus americanus*) population, which instead harbored alternative tetracycline resistance elements (24). With only one exception, human clinical ST1039 strains also harbored Tn*916.1039*.

Interestingly, the Tn*916.1039*-free SL1039 strain, PNUSAL003269, was phylogenetically distanced from other SL1039 (Fig. 1; Fig. S1). This may suggest a bifurcation event in the evolution of SL1039, with one branch containing the Tn*916.1039*-free strain and the other including all other SL1039 strains that may have become amplified and disseminated upon acquisition of the transposon. This hypothesis is supported by the conserved location of this transposon in SL1039 reported previously (24) and confirmed with the current findings (Table 1). As previously speculated (24), the 38% GC content of Tn*916.1039* suggests horizontal gene transfer (HGT) from a microbe with GC content similar to *L. monocytogenes* (e.g., *Enterococcus*). HGT events can be major drivers for the evolution of novel clones in bacterial pathogens (41–44). To our knowledge, SL1039 is the only known *L. monocytogenes* clone for which tetracycline resistance is an almost universally conserved clonal feature. The potential contributions of Tn*916.1039* in the adaptive physiology, virulence and ecology of *L. monocytogenes* remain to be elucidated. Preliminary findings suggest markedly reduced survival in water of the Tn*916.1039*-free strain PNUSAL003269 in comparison to other SL1039 strains (A. Sadat, P. Brown, S. Kathariou, unpub. data).

The type II RM system LmoJ3 was detected in all SL782 and SL1039 strains in a chromosomal hotspot (*lmo0301*-*lmo0305*) for diverse RM systems (45–48). SL782 and SL1039 exhibited an unusual resistance profile with wide-host-range phages: they were susceptible to phages A511 and P100, isolated in 1990 and 1997 from sewage in Germany (26,27) but resistant to phages 20422-1 and 805405-1, isolated in 2004 and 2005 from food processing environments in the USA (25). It is conceivable that 20422-1 and 805405-1 may be also commonly encountered in North American natural environments such as water and wildlife, and resistance to these phages promotes persistence of SL782 and SL1039 in such environmental reservoirs.

Conserved clone-wide phage resistance in *L. monocytogenes* is uncommon, only reported in another relatively recent clone, CC6 (formerly “epidemic clone II”), for which the thermoregulated type II RM system LmoH7 constitutes a clone-specific feature (47).

A surprising trait of SL782 and SL1039 is their geographical distribution. To date the strains have derived exclusively from North America, specifically the United States. It is thus noteworthy that SL2 is currently the only CC2 sublineage with global distribution, and all five remaining sublineages within this clonal complex (SL781, SL782, SL1039, SL1787 and SL2377) are exclusively of North American origin. It is furthermore noteworthy that SL1039 shows strong proclivity for the Eastern Seaboard region of the United States. This may reflect the distribution of specific wildlife hosts or other reservoirs, wildlife mobility patterns, or potentially the movement of specific human ethnic groups and their preferred food products along the Eastern Seaboard. We expect that SL782 and SL1039 may eventually disseminate from North America to other continents, e.g., via international travel and the global food trade, similarly to the major *L. monocytogenes* clone CC6, which was first detected in the 1990s in North America but has since disseminated worldwide (12,15,21,49).

CC2 appears to be a major, ubiquitous, and ancient clone of *L. monocytogenes* (10). It accounted for a substantial fraction (16/85, ca. 19%) of an international panel of historical (1926–1964) *L. monocytogenes* from various sources and geographical regions, with all of these CC2 strains being SL2, primarily ST2 (16,17). ST2 and closely-related STs in SL2 were implicated in some of the first investigated foodborne listeriosis outbreaks, e.g., in Boston, USA in 1979, 1983 and 1985 and in the United Kingdom in 1989 (18–21). SL2 and specifically ST2 continue to dominate CC2 (12).

SL782 and SL1039 have been newly identified, with the earliest strain being from 2002. It is possible that SL782 and 1039 emerged relatively recently within CC2. Alternatively, they may represent ancient clones that evolved in specialized and until recently under-surveyed reservoirs, e.g., water and wildlife, with limited contact with humans and the human food supply. Intriguing preliminary data suggest that SL782 and SL1039 emerged within CC2 approx. 250-350 years ago (E. Gadin, P. Perot, M. Lecuit, unpublished data). The unique combinations of genetic traits discussed here may have driven the emergence and subsequent amplification of these clones. For SL782, such trait combinations may have included the presence of LIPI-4 and LmoJ3, together with loss of *LMOf2365_1900*, while SL1039-relevant traits would include the presence of LIPI-3 and LmoJ3, along with the acquisition of Tn*916.1039*.

To date, the remaining CC2 sublineages (SL781, SL1787, SL2377) remain rare, each consisting of three or fewer strains. Thus, there are striking differences among CC2 sublineages in their apparent epidemiological success as human foodborne pathogens, with SL2 by far the most successful sublineage in this context. First isolated from human listeriosis patients almost 100 years ago (16,17), it still constitutes the vast majority of CC2 isolates. Such pronounced dominance of a single sublineage was also noted with the two other leading hypervirulent clonal complexes, CC1 and CC6, which currently appear to consist of a single sublineage each (11). In fact, of the four leading hypervirulent clonal complexes of *L. monocytogenes* only CC4 avoids extreme dominance by one sublineage, with SL4 being the majority sublineage while SL219 accounts for the remaining 25% of the strains (11). The potential adaptive physiology or virulence traits responsible for such pronounced differences within CC2 and indeed other leading hypervirulent CCs of *L. monocytogenes* remain unknown and clearly in need for further investigation.

## AUTHOR CONTRIBUTIONS

PB and SK: conceptualization. PB and SK: formal analysis and writing—original draft. SK: funding acquisition. PB, ZK, PP, AS, JJ, DE, EG, ML and SK: investigation. SK: project administration. PB, PP and SK: visualization. PB, ZK, PP, AS, JJ, DE, EG, ML and SK: writing—review and editing. All authors have read and agreed to the published version of the manuscript.

## ACKNOWLEDGEMENTS

We would like to thank Alexandra Moura for her sequencing data gathering and curation contributions as the PubMLST BIGSdb administrator.

## FUNDING STATEMENT

This work was partially supported by a National Alliance for Food Safety and Security grant (USDA) and by award 2018-07464 from the USDA National Institute of Food and Agriculture. Any opinions, findings, conclusions, or recommendations expressed are those of the author and do not necessarily reflect the view of the USDA.

**Disclaimer:** The opinions expressed by authors contributing to this journal do not necessarily reflect the opinions of the Centers for Disease Control and Prevention or the institutions with which the authors are affiliated.

## SUPPLEMENTARY MATERIALS

**Supplementary Table 1.** *Listeria monocytogenes* CC2 strains employed in phylogenetic tree construction (Fig. 1).

**Supplementary Table 2.** Core genome multilocus sequence typing (cgMLST) distance matrix between strains shown in Fig. S1. Each number represents the number of cgMLST allele differences between the two respective strains. Strain IDs and BIGSdb Isolate IDs are provided for each strain. Strains of SL2, SL782 and SL 1039 are highlighted in green, blue and red, respectively. Included strains were chosen from the Centers for Disease Control & Prevention database and from publications with publicly available sequence data for CC2 strains (5,6,17,48).

**Supplementary Table 3.** Additional genomic feature information from CC2 strains found in Table 1. Features selected from Maury et al. 2016 and Wiktorczyk-Kapischke et al. 2023 (12,32).

**Supplementary Figure 1.** Phylogeny of clonal complex (CC) 2. The analyzed genomes included representative CC2 strains listed in Table 1, previously investigated using a cgMLST-based approach (50) and other CC2 strains from the Centers for Disease Control & Prevention database. The phylogenetic tree was created using a cgMLST-based approach as described in Materials and Methods. Scale bar represents the percent identity of the 1,748 cgMLST alleles with cgMLST differences ranging from 0 to 1,174 alleles. Sequence type (ST), sublineage (SL) and cgMLST type (CT) designations are provided for each strain.

