## Supplementary figures and images for "Novel, highly divergent clones in *Listeria monocytogenes* serotype 4b in North America: Sublineages 782 and 1039, members of the hypervirulent clonal complex 2"

### Supplemental Figure 1

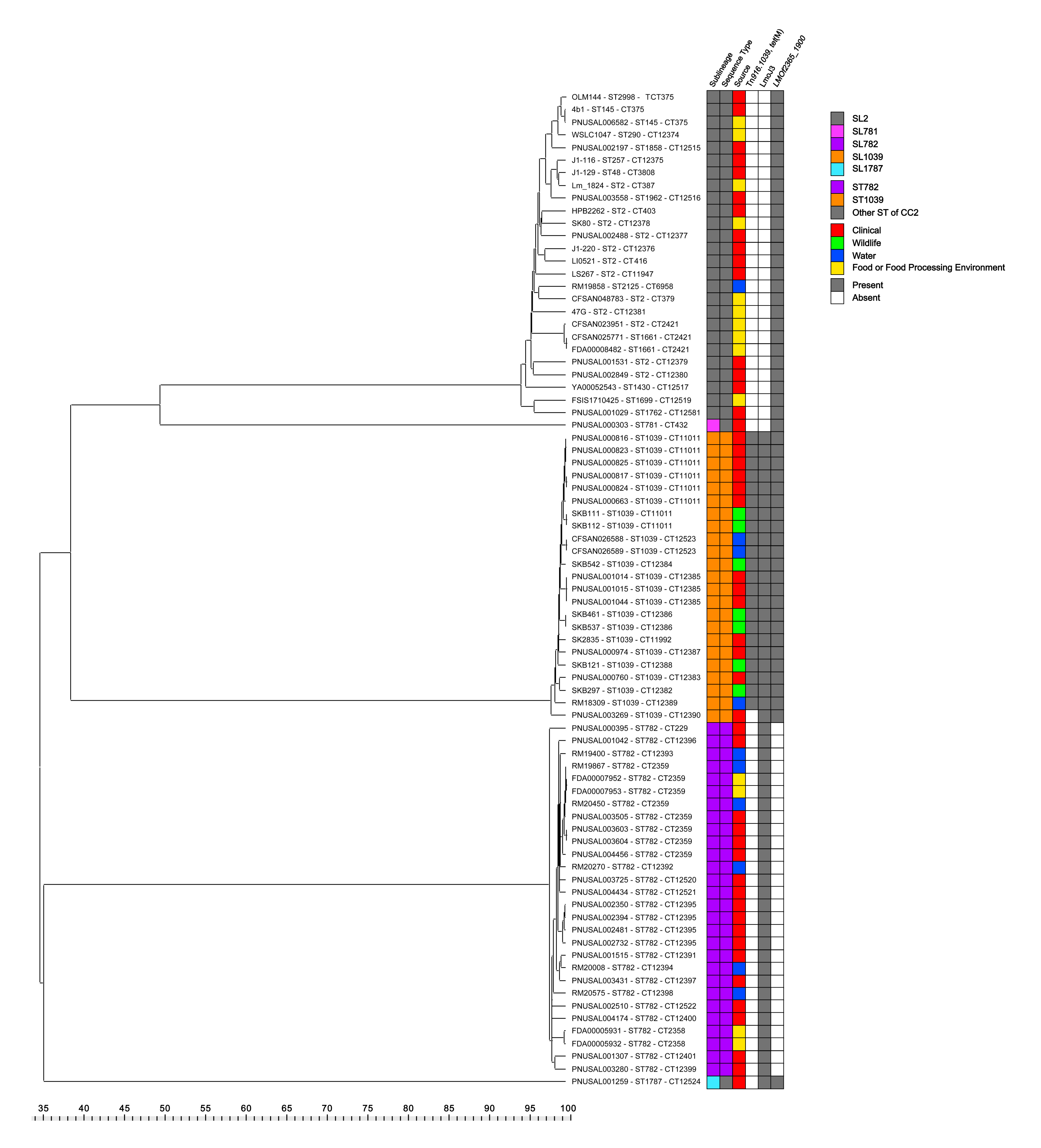
